# Comparison of Clustering Methods for High-Dimensional Single-Cell Flow and Mass Cytometry Data

**DOI:** 10.1101/047613

**Authors:** Lukas M. Weber, Mark D. Robinson

## Abstract

Recent technological developments in high-dimensional flow cytometry and mass cytometry (CyTOF) have made it possible to detect expression levels of dozens of protein markers in thousands of cells per second, allowing cell populations to be characterized in unprecedented detail. Traditional data analysis by “manual gating” can be inefficient and unreliable in these high-dimensional settings, which has led to the development of a large number of automated analysis methods. Methods designed for unsupervised analysis use specialized clustering algorithms to detect and define cell populations for further downstream analysis. Here, we have performed an up-to-date, extensible performance comparison of clustering methods for high-dimensional flow and mass cytometry data. We evaluated methods using several publicly available data sets from experiments in immunology, containing both major and rare cell populations, with cell population identities from expert manual gating as the reference standard. Several methods performed well, including `FlowSOM, X-shift, PhenoGraph, Rclusterpp`, and `flowMeans`. Among these, `FlowSOM` had extremely fast runtimes, making this method well-suited for interactive, exploratory analysis of large, high-dimensional data sets on a standard laptop or desktop computer. These results extend previously published comparisons by focusing on high-dimensional data and including new methods developed for CyTOF data. R scripts to reproduce all analyses are available from GitHub (https://github.com/lmweber/cytometry-clustering-comparison), and pre-processed data files are available from FlowRepository (FR-FCM-ZZPH), allowing our comparisons to be extended to include new clustering methods and reference data sets.

## Introduction

Flow cytometry is a widely used technology for identifying and quantifying cell types (populations) by measuring expression levels of surface and intracellular proteins in individual cells. In immunology, experimental settings include: detecting specific cell populations such as disease biomarkers; characterizing unknown cell populations; and quantifying differences in population abundance between samples in different conditions, such as diseased and healthy. Modern flow cytometers can routinely detect 15-20 parameters (protein markers) per cell [1,2], at throughput rates above 10,000 cells per second. State-of-the-art systems may reach as many as 50 parameters [2]. Detecting a large number of parameters per cell allows populations to be characterized in great detail. However, the number of parameters is ultimately limited by technical issues such as spectral overlap and autofluorescence [3].

Mass cytometry (also known as CyTOF, for “cytometry by time-of-flight”) is a recent technological development [4]. Instead of using fluorescent tags, antibodies are labeled with transition metal isotopes, and antibody-stained cells are passed through a time-of-flight mass spectrometer. By using metal isotopes instead of fluorescent tags, mass cytometry greatly reduces the problem of spectral overlap and eliminates autofluorescence, resulting in the ability to detect a greater number of parameters per cell. Currently, mass cytometry systems can typically measure around 40 parameters per cell, and this could theoretically increase to more than 100 [4]. Throughput rates are on the order of hundreds of cells per second. Unlike flow cytometry, it is not possible to collect cells after an experiment, as they are destroyed during the mass spectrometry step.

Data analysis for flow cytometry has traditionally been done by “manual gating”, which consists of visual inspection of two-dimensional scatterplots to identify known cell populations. However, this technique suffers from several major limitations, including subjectivity, operator bias, difficulties in detecting unknown cell populations, and difficulties in reproducibility [2, 5, 6]. These problems are especially pronounced in high-dimensional settings (large numbers of parameters per cell), since there are too many two-dimensional projections to reliably analyze, and any multidimensional structure not seen in the twodimensional projections is ignored. To address these issues, major efforts have been made to develop partially or fully automated analysis methods.

Automated analysis methods may be grouped into two main categories: unsupervised and supervised [2]. Unsupervised approaches use clustering methods to detect cell populations, defined here as groups of cells with similar protein marker expression profiles. Clustering analysis may be performed on data from a single biological sample, on data from multiple samples on a per-sample basis, or on combined data from multiple samples. Detected clusters (cell populations) can then be analyzed individually or compared across samples, for example by comparing cluster frequencies between samples in different biological conditions. Importantly, this procedure allows previously unknown cell populations to be described in an unbiased, data-driven manner; this type of exploratory analysis is difficult or impossible with manual gating, especially when using high-dimensional data.

By contrast, supervised approaches rely on an external biological or clinical variable describing each sample. This could be a simple categorical variable such as disease status or tissue type, or a more complex clinical outcome such as survival time. Supervised approaches use this external variable as an input to train a model, which can then be used to predict the status of new samples. Many supervised approaches will also return an interpretable model; for example returning a set of cell populations correlated with the external variable, which may be investigated as possible biomarkers.

During the last 5-10 years, many new automated analysis methods have been proposed, but guidance for researchers and bioinformaticians interested in applying them has been difficult to find. To address this, the FlowCAP (“Flow Cytometry: Critical Assessment of Population Identification Methods”) Consortium organized a series of challenges to objectively evaluate the performance of the various methods, using standardized benchmark data sets. The FlowCAP-I challenges evaluated unsupervised methods, finding that several automated methods were able to accurately reproduce expert manual gating [7]. Subsequent FlowCAP challenges focused on supervised approaches. The FlowCAP-IV challenge used a complex data set containing a clinical survival time variable for samples from a large number of patients, and found that two methods were able to generate statistically significant predictive value [8]. FlowCAP-III (challenge 4) also tested an intermediate “semi-supervised” approach, where the cell population hierarchy was provided; automated methods were able to match the performance of centralized manual gating for a large, multi-laboratory data set [6].

In this study, we focus on unsupervised approaches. While supervised approaches may be superior when external status or outcome variables are available across multiple biological samples, there are many situations where these variables do not exist. In particular, unsupervised approaches can be used for exploratory analysis, for example to investigate the diversity of cell populations within a single sample; this type of exploratory analysis is not possible in a supervised context. Several recent studies have also used unsupervised approaches to compare frequencies of detected cell populations between groups of samples in different biological conditions, using high-dimensional CyTOF data [9–11]. However, the FlowCAP-I challenges did not include any high-dimensional benchmark data sets, making it difficult to interpret the FlowCAP-I findings for new studies involving CyTOF data. Due to the “curse of dimensionality”, the performance of clustering algorithms in low-dimensional settings is in general not a good guide to performance in higher-dimensional settings [12, 13]; both clustering accuracy and computational efficiency may be severely affected, depending on the mathematical properties of the algorithm.

In addition, since the publication of the FlowCAP-I results, several new clustering methods designed specifically for CyTOF data have been published. A number of recent studies have provided overviews of available clustering methods for high-dimensional cytometry data [1, 2, 14–16], performance comparisons against a subset of existing methods while introducing a new method [17,18], or performance comparisons using simulated data [19]. However, a comprehensive, updated benchmarking of methods using high-dimensional experimental data sets has been lacking.

In this study, we have performed an up-to-date, extensible performance comparison of clustering methods for high-dimensional flow and mass cytometry data. This includes several new methods that were not yet available at the time of the FlowCAP-I challenges, and which have been developed specifically with high-dimensional CyTOF data in mind. Unlike FlowCAP-I, which used data sets with low to moderate dimensionality, we have used highdimensional data sets, since clustering algorithms may behave very differently in these settings. We have selected six publicly available data sets, where cell population identities are known from expert manual gating. The data sets contain major and rare immune cell populations in well-characterized biological systems, where manual gating is likely to be reliable despite the high dimensionality. We test two distinct clustering tasks: detection of all major immune cell populations, and detection of a single rare cell population of interest. The clustering methods are evaluated by their ability to reproduce the expert manual gating, using an extension of the original FlowCAP-I methodology. Our aim is to provide guidance to researchers and bioinformaticians interested in applying clustering methods for unsupervised analysis of new data sets from experiments in high-dimensional cytometry. Code and pre-processed data files are available from GitHub (https://github.com/lmweber/cytometry-clustering-comparison) and FlowRepository (repository FR-FCM-ZZPH), allowing our analyses to be easily reproduced or extended to include new clustering methods and reference data sets.

## Materials and Methods

### Clustering methods

We compared a total of 18 clustering methods (Table 1). All of these methods are freely available; we did not include any methods without freely available software implementations, since our aim is to provide practical recommendations to researchers performing data analyses. Our results also do not include methods that we were unable to run successfully (Supplementary Methods). The methods are based on a wide range of theoretical approaches, which are briefly described in Table 1. For detailed explanations of the approaches, we refer to the original references. Software package versions are listed in Supplementary Table S1. In addition to the individual clustering analysis, we also performed ensemble clustering (consensus clustering) using the clue R package [20] (Supplementary Methods), as done previously in the FlowCAP-I challenges.

**Table 1.**
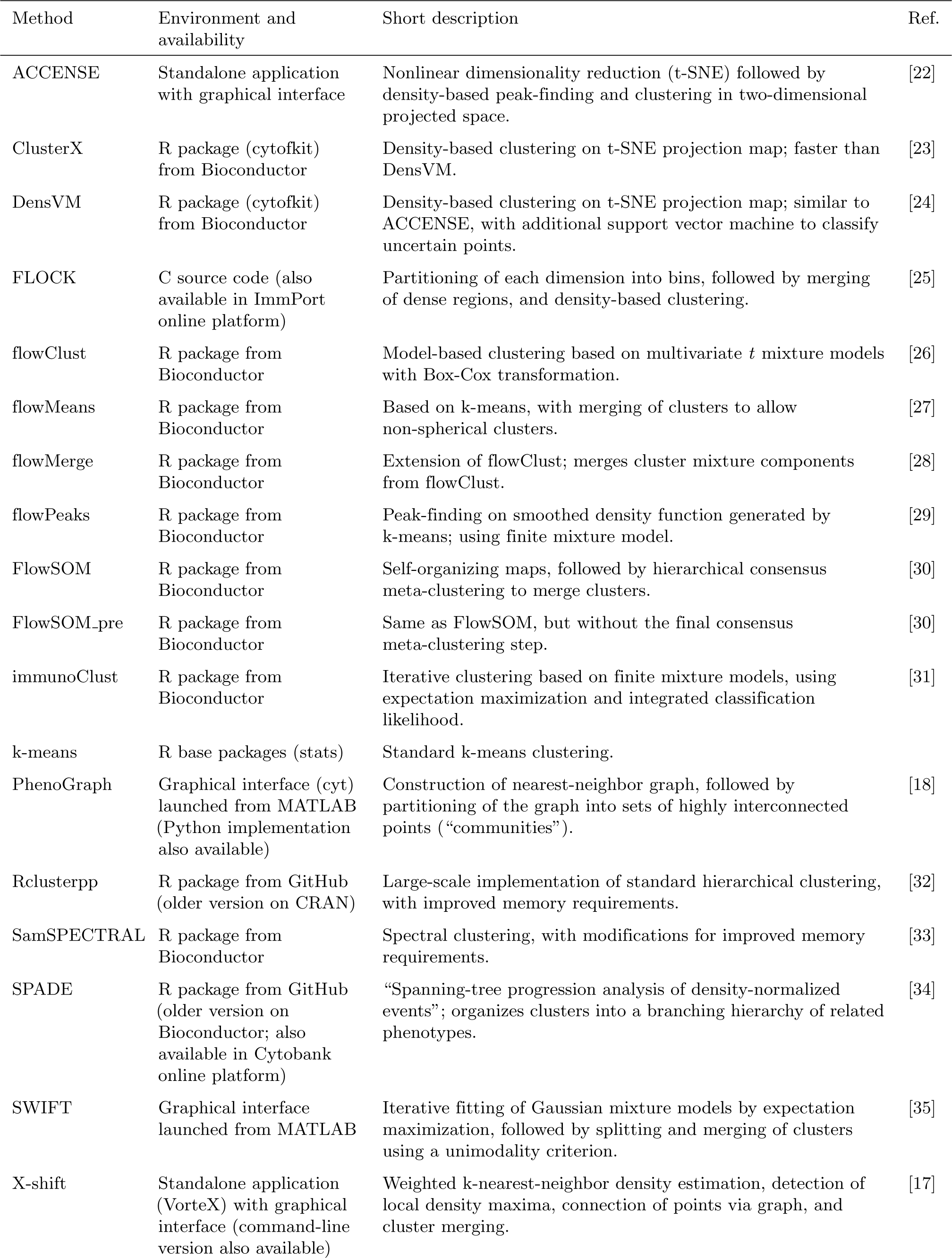
Overview of clustering methods compared in this study.

Additional details including software package versions and parameter settings used for each clustering method are included in Supplementary Table S1.

### Data sets

To evaluate the clustering methods, we selected six publicly available data sets from experiments in immunology using CyTOF or high-dimensional flow cytometry (Table 2). Throughout the comparisons, we use manually gated cell population labels as the reference populations, or “ground truth”, against which the clustering algorithms are evaluated; the data sets are from well-characterized biological systems, where manual gating is likely to be reliable even in high-dimensional settings. For each of these data sets, manually gated population labels are available either directly within the data files published by the original authors, or are reproducible from published gating diagrams.

**Table 2.** Summary of data sets used to evaluate clustering methods.

| Data set | CyTOF or flow cytometry | Clustering task | No. of cells | No. of dimensions | No. of manually gated populations of interest | No. of manually gated cells | Organism | No. of individuals (patients, mice) | Sample description | Ref. |
| --- | --- | --- | --- | --- | --- | --- | --- | --- | --- | --- |
| Levine_32dim | CyTOF | Multiple populations | 265,627 | 32 (surface markers) | 14 | 104,184 (39%) | Human | 2 | Bone marrow cells from healthy donors | [18] |
| Levine_13dim | CyTOF | Multiple populations | 167,044 | 13 (surface markers) | 24 | 81,747 (49%) | Human | 1 | Bone marrow cells from healthy donor | [18] |
| Samusik_01 | CyTOF | Multiple populations | 86,864 | 39 (surface markers) | 24 | 53,173 (61%) | Mouse | 1 | Replicate bone marrow samples from C57BL/6J mice (sample 01 only) | [17] |
| Samusik_all | CyTOF | Multiple populations | 841,644 | 39 (surface markers) | 24 | 514,386 (61%) | Mouse | 10 | Replicate bone marrow samples from C57BL/6J mice (all samples) | [17] |
| Nilsson_rare | Flow cytometry | Rare population | 44,140 | 13 (surface markers) | 1 (hematopoietic stem cells) | 358 (0.8%) | Human | 1 | Bone marrow cells from healthy donor | [36] |
| Mosmann_rare | Flow cytometry | Rare population | 396,460 | 14 (surface and intracellular) | 1 (activated memory CD4 T cells) | 109 (0.03%) | Human | 1 | Peripheral blood cells from healthy donor, stimulated with influenza antigens | [35] |

We ran clustering methods on all cells (including unassigned cells, i.e. those not assigned to any population by manual gating), and evaluated performance on the cells where manually gated population labels were available (Table 2). The data sets contain both major and rare cell populations, allowing us to test performance on two distinct clustering tasks: detection of all major immune cell populations, and detection of a single rare cell population of interest.

In order to compare against the previous FlowCAP results, we also included the two highest-dimensional data sets from FlowCAP-I (Supplementary Table S2). However, these data sets are still relatively low-dimensional compared to the six main data sets. Therefore, the results from the main data sets should be used for inferring performance on new high-dimensional data sets.

Pre-processed data files are available for download from FlowRepository (repository FR-FCM-ZZPH) [21]. Original data files can be obtained through the references in Table 2.

### Data pre-processing and parameter settings

Data pre-processing included the application of an arcsinh transformation with a standard cofactor of 5 (CyTOF data) or 150 (flow cytometry data) [4], i.e. *arcsinh*(*x/*5) or *arcsinh*(*x/*150). For the flow cytometry data sets, pre-gating to exclude doublets, debris, and dead cells was also required (Supplementary Methods and Supplementary Figures S37–S38). The clustering algorithms were run on all remaining single, live cells; no additional pre-gating was performed, since our aim is to evaluate performance in maximally automated settings. In addition, we did not perform any standardization of individual protein marker dimensions. This was unnecessary since the arcsinh already transforms all dimensions to comparable scales; and importantly, standardization of dimensions that do not contain a true signal could amplify the effect of noise and outliers, adversely affecting clustering performance.

For each clustering method, we experimented with input parameters in order to give the best possible performance. For many methods, the most important input parameters related to the number of clusters. Some methods provided an option to select the number of clusters automatically; some methods allowed the user to adjust the number indirectly through other parameters; and some methods left it as a direct user input (Supplementary Table S3). We used the automatic option where this was available and gave reasonable results, and otherwise selected 40 clusters for each data set, or adjusted indirect parameters to get close to 40 clusters. The choice of 40 clusters was designed to be conservative, in the sense of tending to select too many clusters rather than too few, in order to avoid smaller populations merging into larger ones (see Supplementary Methods and Supplementary Figure S21 for more details). Supplementary Table S3 summarizes the final number of clusters for each clustering method and data set. All other parameter settings used in the final results are recorded in Supplementary Table S1.

### Evaluation methodology

Our evaluation strategy was largely based on the FlowCAP-I methodology. As in FlowCAP-I, we used the F1 score (harmonic mean of precision and recall) as our main evaluation criterion. The F1 score provides a value between 0 and 1 for each cluster, with 1 indicating a perfect reproduction of the corresponding manually gated population. High precision implies a low proportion of false positives, and high recall (sensitivity) implies low false negatives.

However, we made two important changes to the methodology. Firstly, we matched clusters to reference populations (manually gated populations) using the Hungarian assignment algorithm, which solves the assignment problem by finding a one-to-one mapping that maximizes the sum of F1 scores across reference populations. The use of the Hungarian algorithm in this context was recently introduced by [17]. By contrast, in FlowCAP-I, clusters were matched to reference populations by maximizing the F1 score individually for each population, which potentially allows the same cluster to map to multiple reference populations. For data sets with only a single population of interest (see Table 2), we selected the cluster maximizing the F1 score for this population, since there is no ambiguity in this case.

Secondly, after mapping clusters to reference populations, we used unweighted averages to calculate the mean precision, mean recall, and mean F1 score across reference populations. We used unweighted averages in order to give equal representation to both large and small populations. By contrast, FlowCAP-I used averages weighted by population size, which gives more importance to relatively larger populations. For data sets with only a single population of interest, we reported the precision, recall, and F1 score for this population.

We also recorded runtimes, since these varied by several orders of magnitude between the various methods. While the runtimes are not precisely comparable between methods due to differences in subsampling, number of processor cores, and hardware specifications (Supplementary Tables S1 and S4), the order-of-magnitude differences provide important information for users.

To further investigate the quality of the clustering results, we examined the protein expression profiles of detected clusters, and compared them against reference populations using heatmaps together with hierarchical clustering. Finally, we investigated stability of the clustering results by running methods multiple times with different random starts and bootstrap resamples (Supplementary Methods).

## Results

### Detection of multiple cell populations

The results of the performance comparison are summarized in Table 3. The first four data sets (`Levine_32dim, Levine_13dim, Samusik_01`, and `Samusik_all`) contain multiple cell populations of interest (see Table 2). For these data sets, the mean F1 score across reference populations is shown; the best-performing methods are `FlowSOM` (data sets `Levine_32dim, Samusik_01, and Samusik_all`) and `flowMeans` (data set `Levine_13dim`). Several other methods also consistently perform well, including `X-shift, PhenoGraph, ClusterX, FLOCK`, and `Rclusterpp`.

**Table 3.** Results of comparison of clustering methods.

|  | Multiple populations of interest |  |  |  |  |  |  |  | Single rare population of interest |  |  |  |
| --- | --- | --- | --- | --- | --- | --- | --- | --- | --- | --- | --- | --- |
|  | Levine_32dim |  | Levine_13dim |  | Samusik_01 |  | Samusik_all |  | Nilsson_rare |  | Mosmann_rare |  |
|  | mean F1 | runtime<br>hh:mm:ss | mean F1 | runtime<br>hh:mm:ss | mean F1 | runtime<br>hh:mm:ss | mean F1 | runtime<br>hh:mm:ss | F1 | runtime<br>hh:mm:ss | F1 | runtime<br>hh:mm:ss |
| ACCENSE | 0.494 | 00:05:32 | 0.358 | 00:04:48 | 0.517 | 00:06:21 | 0.502 | 00:05:32 | 0.445 | 00:06:11 | 0.021 | 00:04:37 |
| ClusterX | **0.682** | **01:57:02** | **0.474** | **03:50:51** | 0.571 | 01:52:09 | 0.603 | 02:02:08 | 0.132 | 00:29:00 | 0.004 | 01:56:13 |
| DensVM | 0.660 | 08:30:13 | 0.448 | 08:11:09 | 0.239 | 07:34:49 | 0.496 | 07:55:14 | 0.153 | 03:19:36 | 0.004 | 07:55:34 |
| FLOCK | **0.727** | **00:03:43** | 0.379 | 00:00:29 | 0.608 | 00:00:35 | **0.631** | **00:14:28** | 0.089 | 00:00:08 | 0.102 | 00:01:06 |
| flowClust | NA | NA | 0.416 | 02:59:27 | 0.612 | 06:04:13 | 0.610 | 11:56:58 | **0.461** | **04:20:24** | 0.080 | 03:32:41 |
| flowMeans | **0.769** | **02:34:01** | **0.518\*** | **00:04:09** | 0.625 | 04:13:12 | **0.653** | **02:03:17** | **0.488** | **00:01:06** | 0.104 | 00:03:57 |
| flowMerge | NA | NA | 0.247 | 07:45:41 | 0.452 | 09:56:25 | 0.341 | 03:21:40 | 0.111 | 09:41:02 | 0.159 | 11:06:45 |
| flowPeaks | 0.237 | 00:05:19 | 0.215 | 00:00:21 | 0.058 | 00:07:05 | 0.323 | 00:16:39 | 0.016 | 00:00:08 | 0.001 | 00:02:18 |
| FlowSOM | **0.780\*** | **00:00:41** | **0.495** | **00:00:15** | **0.707\*** | **00:00:19** | **0.702\*** | **00:02:13** | **0.447** | **00:00:08** | **0.665** | **00:02:14** |
| FlowSOM_pre | 0.502 | 00:00:35 | 0.422 | 00:00:10 | 0.583 | 00:00:14 | 0.528 | 00:02:08 | **0.447** | **00:00:03** | **0.665** | **00:01:32** |
| immunoClust | 0.413 | 03:20:51 | 0.308 | 02:57:27 | 0.552 | 01:35:10 | 0.523 | 02:06:40 | 0.371 | 00:06:57 | 0.563 | 01:51:23 |
| k-means | 0.420 | 00:00:13 | 0.435 | 00:00:02 | **0.650** | **00:00:05** | 0.590 | 00:00:26 | 0.243 | 00:00:01 | 0.103 | 00:00:11 |
| PhenoGraph | 0.563 | 00:37:00 | **0.468** | **00:12:09** | **0.671** | **00:05:55** | **0.653** | **05:30:35** | 0.229 | 00:01:58 | 0.498 | 00:43:43 |
| Rclusterpp | 0.605 | 01:13:04 | 0.465 | 00:17:54 | **0.637** | **00:08:32** | 0.613 | 00:14:05 | 0.360 | 00:00:17 | **0.737** | **02:12:32** |
| SamSPECTRAL | 0.512 | 04:24:05 | 0.253 | 00:24:01 | 0.263 | 00:34:42 | 0.138 | 00:39:26 | 0.088 | 00:01:52 | **0.618** | **03:42:28** |
| SPADE | NA | NA | 0.127 | 00:04:46 | 0.169 | 00:03:02 | 0.130 | 00:53:39 | 0.180 | 00:00:52 | 0.027 | 00:12:12 |
| SWIFT | 0.177 | 02:27:39 | 0.179 | 01:07:03 | 0.202 | 02:19:30 | 0.208 | 02:50:08 | 0.390 | 00:11:26 | 0.484 | 00:34:34 |
| X-shift | **0.691** | **04:45:26** | **0.470** | **00:48:17** | **0.679** | **00:24:54** | **0.657** | **03:48:27** | **0.531\*** | **00:04:37** | **0.802\*** | **03:18:20** |

Results show the mean F1 score for data sets with multiple cell populations of interest, and F1 score for data sets with a single rare cell population of interest; as well as runtimes. For each data set, the best-performing method is indicated with a star (*), and the top five methods are displayed in bold. Runtimes are not precisely comparable between methods due to differences in subsampling, number of processor cores, and hardware specifications (Supplementary Tables S1 and S4); however they are included in order to provide users with information about order-of-magnitude differences. NA = not available, due to errors or non-completion (Supplementary Table S1).

Among these high-performing methods, `FlowSOM` has by far the fastest runtimes, followed by `FLOCK`. For example, for the largest data set, `Samusik_all` (841,644 cells and 39 dimensions; Table 2), `FlowSOM` ran in less than 3 minutes, without any subsampling required. By contrast, `PhenoGraph` took more than 5 hours, and `X-shift` took more than 3 hours with subsampling to 300,000 cells (Table 3; Supplementary Tables S1 and S4).

Figure 1 provides more detailed results for the first data set, `Levine_32dim`. Clustering performance varies widely between methods in terms of mean F1 score, mean precision, and mean recall (panels A–C). Performance also varies between the individual reference populations; most methods show poor performance for at least one individual population (panel B). These tend to be the relatively smaller populations (panel D and Supplementary Figures S7–S10). Runtimes vary across several orders of magnitude (panel E). However, the best-performing method in terms of mean F1 score for this data set (`FlowSOM`) is also one of the fastest (panel F); this observation represents one of the key results from this study.

**Figure 1.**
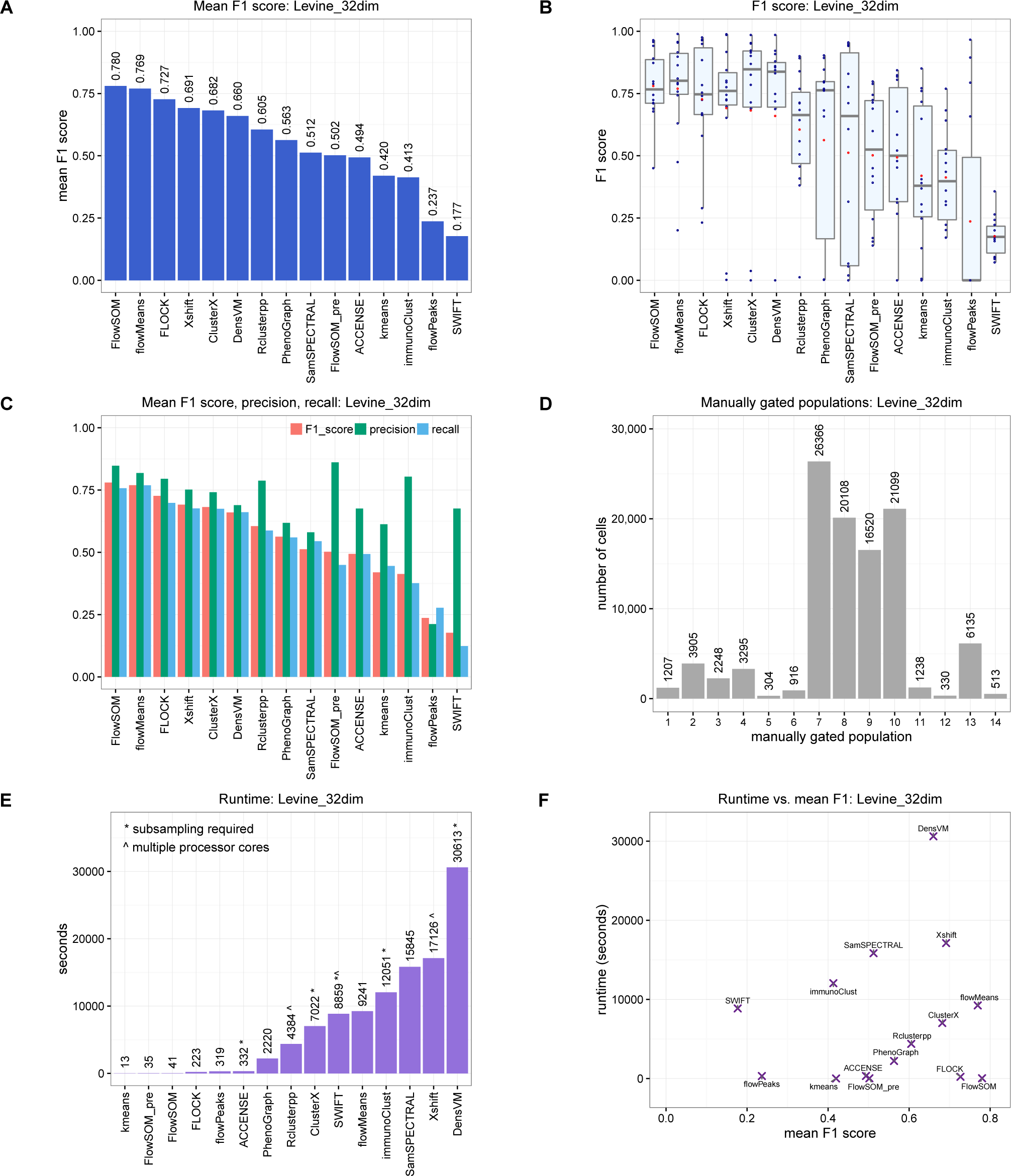
Results of comparison of clustering methods for data set Levine_32dim. (A) Mean F1 score across cell populations. (B) Distributions of F1 scores across cell populations. The box plots show medians, upper and lower quartiles, whiskers extending to 1.5 times the interquartile range, and outliers, with means shown additionally in red. (C) Mean F1 scores, mean precision, and mean recall. (D) Number of cells per reference population. (E) Runtimes. (F) Runtime vs. mean F1 score; methods combining high mean F1 scores with fast runtimes are seen toward the bottom-right. Similar figures of results for all data sets are included in Supplementary Figures S1–S6.

Similar figures of results for the other data sets (`Levine_13dim, Samusik_01`, and `Samusik_all`) are included in Supplementary Figures S2–S4. While the ranking of methods changes somewhat between data sets (see also Table 3), the observation that `FlowSOM` combines best or near-best mean F1 scores with extremely fast runtimes remains consistent.

To investigate the interpretability of the clustering results, we also compared the protein expression profiles of detected clusters against reference populations. Figure 2 shows an example of these results, for `FlowSOM` with data set Levine_32dim. The heatmap displays median expression intensities for each protein marker, with hierarchical clustering to group rows and columns. For most of the reference populations (red rows), at least one detected cluster (blue rows) matches closely, indicating that the clusters correctly correspond to the reference populations. However, the expression profiles do not match perfectly, and some additional splitting of clusters is apparent. Additional figures for all clustering methods for this data set are included in Supplementary Figures S22–S36. Among several of the lower-ranked methods, a significant number of mismatches (red rows grouping together before matching to any blue rows) occur. This demonstrates that the high-performing methods in terms of mean F1 score also generate relatively interpretable clusters.

**Figure 2.**
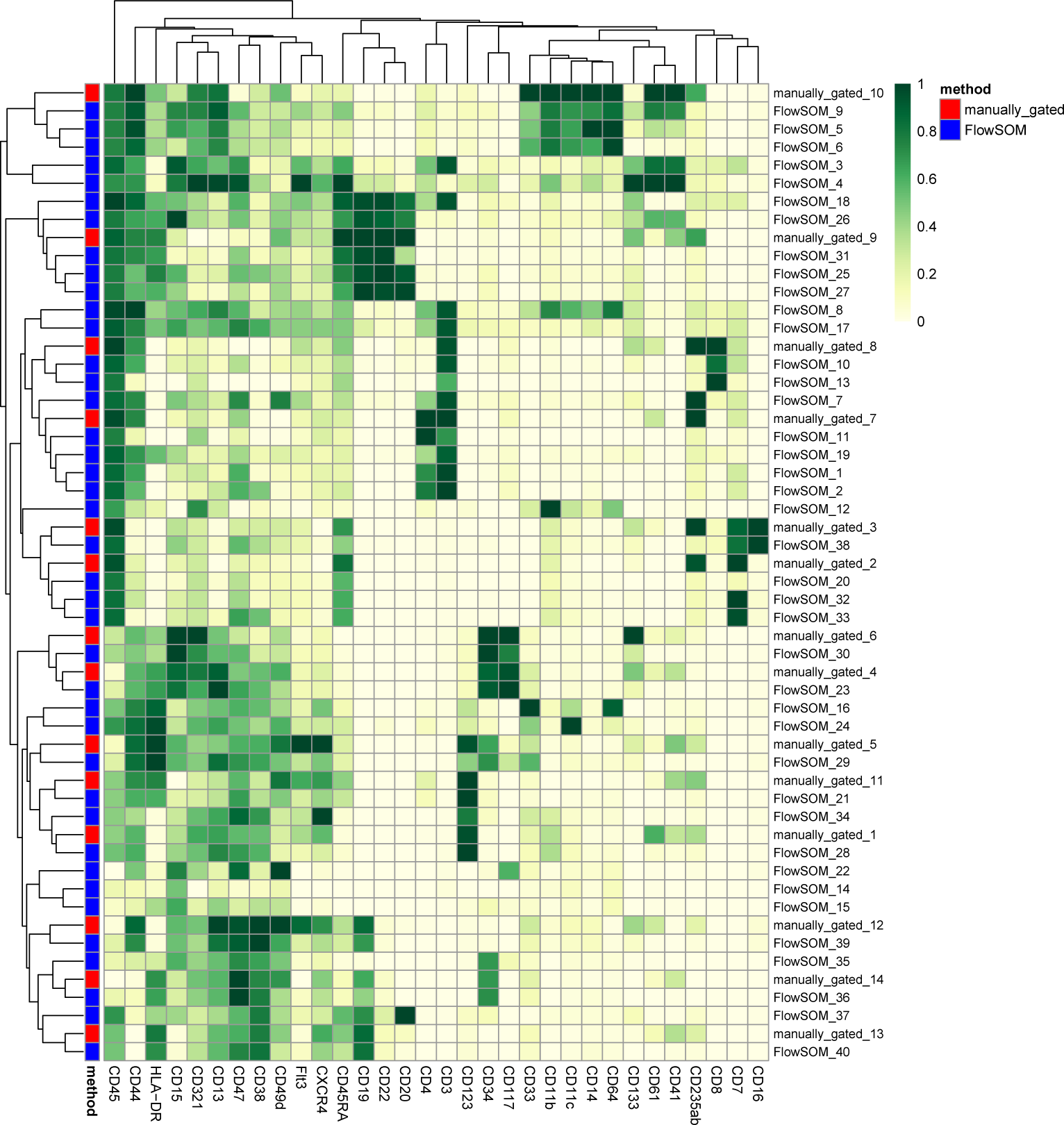
Expression profiles of detected clusters and reference populations, FlowSOM, data set Levine_32dim. Heatmap shows median expression intensities of each protein marker (columns), for each detected cluster or reference population (rows). Values are arcsinh-transformed, and scaled between 0 and 1 for each protein marker. Rows and columns are sorted by hierarchical clustering (Euclidean distance, average linkage). Cluster and population indices are included in row headings. Red labels indicate rows representing reference populations, and blue labels indicate clusters detected by FlowSOM. For most reference populations (red rows), the expression profile of at least one detected cluster (blue rows) matches closely. Additional figures for all clustering methods are included in Supplementary Figures S22–S36.

### Detection of rare cell populations

The last two data sets in Table 3 (`Nilsson_rare` and `Mosmann_rare`) each contain a single rare cell population of interest. `Nilsson_rare` contains a population of hematopoietic stem cells (HSCs), representing 0.8% of total cells; and `Mosmann_rare` contains a population of activated (cytokine-producing) influenza-specific memory CD4 T cells, representing 0.03% of total cells (Table 2).

For these data sets, Table 3 displays the F1 score for the rare population of interest. The best-performing method is `X-shift`, for both data sets. This is followed by `flowMeans` (`Nilsson_rare`) and Rclusterpp (`Mosmann_rare`). `FlowSOM` and `FlowSOM_pre` are within the top five methods for both data sets; as previously, these have by far the fastest runtimes among the top methods. For `Mosmann_rare` (the larger of the two data sets), `FlowSOM` ran in less than 3 minutes, while X-shift required more than 3 hours.

Between these two data sets, the rare population in `Mosmann_rare` represents a much smaller fraction of total cells, suggesting that the clustering task is likely to be more difficult for this data set. Figure 3 provides more detailed results for `Mosmann_rare`. While several methods achieve good results (panel A), more than half of the methods perform poorly (very low F1 scores). For most of the methods with low F1 scores, recall remains high, while precision is low (panel A), implying that these methods were not able to successfully separate the rare population from other, larger populations. `SWIFT` achieved the highest precision, but at the expense of low recall. As previously, runtimes varied widely between methods (panel B). While `X-shift` and `Rclusterpp` achieved the highest F1 scores, `FlowSOM` again combines high F1 scores with extremely fast runtimes (panel C).

**Figure 3.**
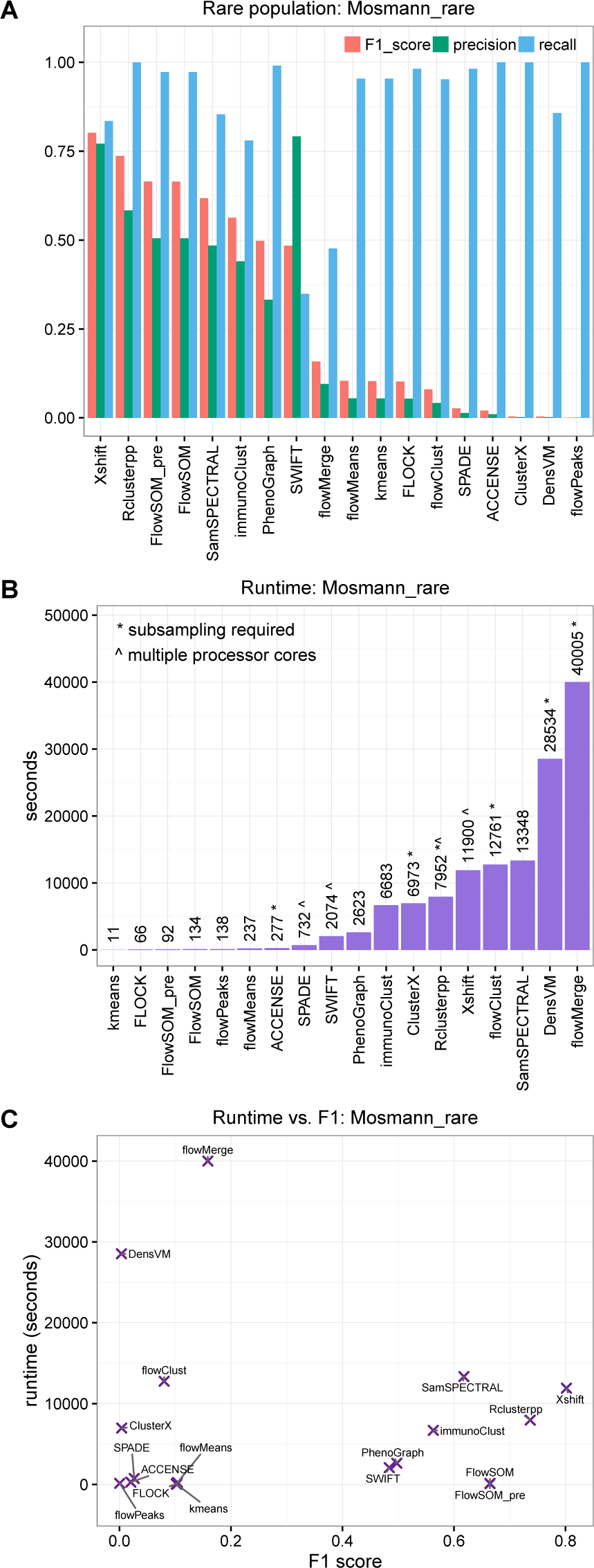
Results of comparison of clustering methods for data set Mosmann_rare. (A) F1 score, precision, and recall for the rare cell population of interest. The rare population contains approximately 0.03% of total cells (Table 2). (B) Runtimes. (C) Runtime vs. F1 score; methods combining high F1 scores with fast runtimes are seen toward the bottom-right. Similar figures of results for all data sets are included in Supplementary Figures S1–S6.

Supplementary Figure S6 displays a similar figure of results for data set `Nilsson_rare`. In addition to `X-shift`, `FlowSOM`, and `FlowSOM_pre`, which again performed well, several other methods that performed poorly for `Mosmann_rare` performed well for `Nilsson_rare` (in particular, `flowMeans`, `flowClust`, and `ACCENSE`). Most methods were also significantly faster, since this is a smaller data set (Tables 2 and 3). Comparing the two data sets, we also observe that `immunoClust`, `SWIFT`, and PhenoGraph performed reasonably well across both data sets, although they were not ranked within either set of top five methods (Table 3).

### FlowCAP data sets

Results for the two highest-dimensional data sets from FlowCAP-I (labeled `FlowCAP ND` and `FlowCAP WNV`; see Supplementary Table S2 for details) are displayed in Supplementary Table S5. Two sets of results are presented, using alternative evaluation methodologies. The first set (first two columns) uses the same methodology as we used for the other data sets in this study, i.e. the Hungarian algorithm to match clusters to reference populations, and unweighted averages to calculate mean F1 scores. The second set (last two columns) uses the original evaluation methodology from FlowCAP-I, i.e. matching clusters by individually maximizing the F1 score for each reference population, and calculating mean F1 scores with weighting by reference population size (see Materials and Methods).

The two sets of results differ significantly. Using the FlowCAP-I methodology, most methods give very high mean F1 scores, consistent with the previously published results from FlowCAP-I [7]. Note that there are some small differences (for those methods available at the time of FlowCAP-I), due to several factors including differences in manually tuned parameters, subsampling, and updated software versions.

By contrast, the mean F1 scores from our updated methodology are lower for most methods. This demonstrates the importance and impact of the choice of evaluation methodology. In our view, the updated methodology is more reliable, since clusters are not allowed to map to multiple reference populations (Hungarian algorithm), and both large and small populations are represented equally (unweighted averages). This avoids the possibility that the mean F1 scores are dominated by one or two large clusters with high individual scores. However, despite the lower scores, several methods still perform well, confirming the main conclusion that automated methods can accurately reproduce expert manual gating for these data sets.

### Ensemble clustering

Results of the ensemble clustering (consensus clustering) are displayed in Supplementary Figures S11–S16. Unlike FlowCAP-I, we found that ensemble clustering did not give any improvements in performance compared to the best-performing individual methods. For the data sets with multiple populations of interest, ensemble clustering gave results similar to the best individual methods. For the data sets with a single rare population of interest, performance was significantly worse; which is surprising. A possible explanation may be that the ensemble clustering performed well for larger populations, but poorly for smaller or rare populations. Due to the change in evaluation methodology (Hungarian algorithm and unweighted averages; see Materials and Methods), the influence of smaller populations has been amplified, hence reducing the overall scores. Further work is warranted in order to investigate these results in more detail.

### Stability of clustering results

The stability analysis (Supplementary Figures S17–S20) revealed that several methods were sensitive to random starts and bootstrap resampling, especially when detecting a single rare cell population. For some methods (such as `FlowSOM`), this included a number of outlier runs, where performance was significantly worse than usual. By contrast, variability was smaller for the data sets with multiple populations of interest (except for `FLOCK`). In each case, the figures display the range of scores recorded over 30 replicate runs per method, with varying random starts or bootstrap resamples. For some methods (`FLOCK`, `flowMeans`, and `flowPeaks`), the bootstrap results are more informative, due to difficulties in accessing internal random seeds for the random starts during parallelized operation (Supplementary Methods).

## Discussion

### Several clustering methods accurately reproduce expert manual gating in highdimensional cytometry data sets

The results showed that several clustering methods were able to accurately detect clusters representing manually gated cell populations in these high-dimensional CyTOF and flow cytometry data sets. In particular, `FlowSOM`, `flowMeans`, `X-shift, PhenoGraph, ClusterX, FLOCK`, and `Rclusterpp` performed well for the data sets with multiple cell populations of interest; and `X-shift`, `FlowSOM`, `FlowSOM_pre`, `Rclusterpp`, `immunoClust`, `SWIFT`, and PhenoGraph performed well for the data sets with a single rare cell population of interest (see Table 3 for full results).

Due to the curse of dimensionality, standard clustering algorithms for low-dimensional data are generally not expected to perform well in high-dimensional settings. Most of the methods tested in this study are specialized clustering algorithms designed for cytometry data. Many of these were published during the last 2–3 years, and are intended for analysis of high-dimensional CyTOF data. The FlowCAP-I comparisons [7] included only lower-dimensional flow cytometry data sets, and many of the methods included here were not yet available at that time; including the two best-performing methods overall, `FlowSOM` and `X-shift`. This study provides an up-to-date comparison, focusing on high-dimensional data sets and including the latest methods for CyTOF data. Our analysis scripts are publicly available, and designed to be extensible in order to accommodate new methods and data sets.

### Runtimes vary widely between methods

Due to the different mathematical and computational approaches taken by the various methods, runtimes varied across several orders of magnitude. An unexpected result from this study was that `FlowSOM`, which gave best or near-best clustering performance for all data sets, also had among the fastest runtimes (Table 3). This demonstrates the importance of choices made during method design with regard to the underlying theoretical clustering approaches; high-performing methods do not necessarily need to be those with the greatest computational requirements.

### Number of clusters

Many methods included options to automatically select the number of clusters, while others left this parameter as a user input or controlled it via indirect parameters (see Materials and Methods, and Supplementary Table S3). An important observation from this study was that the automatic options performed poorly for several methods, and indirect parameters were often difficult to tune. To improve performance, we attempted to select around 40 clusters per data set in these cases (Supplementary Table S3). For example, the automatic option in `FlowSOM` returned too few clusters (less than 10 clusters for each data set; Supplementary Table S1), while `FlowSOM` gave excellent results using 40 clusters per data set (Table 3 and Supplementary Figure S21).

In general, we found that methods providing a simple, direct parameter input to manually adjust the number of clusters were easiest to work with. Among the high-performing methods, this included `FlowSOM`, `flowMeans`, and `Rclusterpp`. Although the number of relevant clusters in an experimental data set may not be known in advance, a direct parameter input allows users to explore the data interactively, for example to find a threshold resolution where a rare population splits from a larger population. This is difficult when the number of clusters is controlled indirectly, and may even be impossible if only an automatic option is presented.

In addition, from a biological point of view, it may be argued that automatically determining the number of clusters is often not a particularly meaningful problem in cytometry data. This is because cell populations may be viewed as effectively forming a near-continuous progression of phenotypes; depending on the desired resolution, an almost arbitrary number of clusters (cell populations) can be defined. (However, some clusters are typically more stable than others, making it difficult to adjust the resolution equally across diverse populations.) Therefore, clustering methods should leave the choice of resolution to the user, so that it may be explored directly.

For many applications, it is generally also “safer” to select slightly too many clusters, rather than too few. During downstream statistical analysis, it is a relatively simple matter to manually merge clusters with similar phenotypes. This conservative strategy helps ensure that smaller or rare populations are adequately separated from larger populations. Providing a simple parameter input for the final number of clusters facilitates this procedure.

### Stability of clustering results

Several methods were sensitive to random starts and bootstrap resampling, especially when detecting a single rare cell population. In particular, we observed a number of outlier runs, where performance was significantly worse than usual (this included `FlowSOM`, the best-performing method overall). By contrast, for the data sets with multiple cell populations of interest, variability was relatively small for most methods. Based on these results, we recommend running clustering methods multiple times with different random starts when the aim is to detect rare cell populations.

### Clustering approaches and alternative analysis procedures

Throughout this study, we have evaluated clustering methods according to their ability to reproduce manual gating in a fully automated manner, with minimal parameter inputs other than the desired number of clusters. However, some of the methods we included are not strictly intended for performing fully automated clustering in this way. For example, the authors of `SWIFT` describe a semiautomated analysis pipeline, where `SWIFT` is initially used to generate a large number of small clusters, and these clusters are then further analyzed by gating (i.e. gating on the clusters). This strategy enables efficient analysis of rare cell populations [35]. Similarly, `immunoClust` is designed to return a relatively large number of clusters [31], some of which may split larger populations. In our evaluations, this negatively affected the reported clustering performance, since our evaluation methodology only allows a single cluster to map to each reference population. We have not attempted to correct for this effect by manually merging clusters, as this would introduce additional subjectivity into the evaluations. Instead, we have compared the ability to perform fully automated clustering, while recognizing that some methods may be more suited for slightly different analysis procedures.

The methods compared in this study are based on a wide variety of theoretical approaches to the clustering problem (Table 1). We have not attempted to judge the relative merits of the theoretical approaches, instead preferring an unbiased empirical evaluation of performance on the chosen experimental data sets. The best-performing method for data sets with multiple cell populations of interest (`FlowSOM`) is based on self-organizing maps and hierarchical consensus meta-clustering, while the top method for detecting a single rare cell population (`X-shift`) employs a completely different strategy based on nearest-neighbor density estimation and graphs (Table 1). Future work could investigate underlying reasons why these approaches perform well.

We also note that many methods required subsampling due to excessive runtimes (Supplementary Tables S1 and S4). This likely had a negative effect on the performance of these methods, especially for the data sets containing rare populations. Depending on the amount of subsampling, rare populations may become difficult to detect if too few cells remain; some algorithms may even exclude them as outliers. In our view, methods that require subsampling for large or high-dimensional data sets are not well-suited for the task of detecting rare populations. In fact, the two best-performing methods (`FlowSOM` and `X-shift`) did not require any subsampling for the data sets where they achieved the best performance respectively (Table 3 and Supplementary Table S4).

### Computational environments and accessibility

The majority of the clustering methods in this study (see Table 1) were available as R packages, most of which were distributed through the Bioconductor project [37]. The remaining methods were available as standalone applications with graphical interfaces, graphical interfaces launched from MATLAB, through online analysis platforms, or as source code (Table 1). The graphical interfaces were designed to be user-friendly and accessible for users without programming experience. However, we found that overall, the methods distributed as R/Bioconductor packages were the easiest to work with. This was due to two main reasons. The first reason related to reproducibility: R packages allow users to write scripts, which can be re-run to generate the same results multiple times (as long as a random seed is specified). This facilitates interactive, exploratory analysis, where users attempt various analyses in an iterative process, with parameter settings recorded in the script. In addition, the reproducibility of final, published results is improved. The second reason related to analysis pipelines and downstream statistical analysis: methods implemented as R packages can be incorporated into full analysis pipelines, from pre-processing to downstream statistical analysis and plotting. The commands for the analysis pipeline are saved in an R script, again facilitating reproducibility, as well as making it easy to add minor adjustments at any point within the pipeline. While graphical interfaces provide more accessibility, we believe this is outweighed by the advantages of the scripting approach. Finally, all Bioconductor packages are distributed with documentation and vignettes, ensuring that users have access to instructions and examples of usage.

### Recommendations

We have performed an up-to-date, extensible performance comparison of clustering methods for automated detection of cell populations during unsupervised analysis of high-dimensional flow and mass cytometry (CyTOF) data. We compared 18 clustering methods, using 6 publicly available data sets from experiments in immunology. Based on our results, we recommend the use of `FlowSOM` (with manual selection of the number of clusters) as a first choice for this type of analysis, since this method gave best or near-best performance across all data sets, together with extremely fast runtimes. Other high-performing methods included `X-shift`, `PhenoGraph`, `Rclusterpp`, and `flowMeans`. Fast runtimes make `FlowSOM` well-suited for performing interactive, exploratory analyses of large data sets (possibly up to millions of cells) on a standard laptop or desktop computer. Several methods (including `FlowSOM`) were sensitive to random starts and bootstrap resampling when detecting rare cell populations; we recommend the use of multiple random starts in these cases. Automatically selecting the number of clusters often did not work well; we found that methods providing a simple parameter input to manually select the final number of clusters were the easiest to work with.

In general, it is advisable to select somewhat more clusters than necessary, as this helps ensure that smaller populations remain adequately separated, and it is a relatively simple matter to manually merge clusters during downstream analysis. Finally, we recommend that researchers run methods via a scripting approach wherever possible (for example using R/Bioconductor packages), to facilitate reproducibility and integration into analysis pipelines and downstream statistical analysis. R scripts and pre-processed data files to reproduce our analyses are available from GitHub (https://github.com/lmweber/cytometry-clustering-comparison) and FlowRepository (repository FR-FCM-ZZPH), allowing our comparisons to be extended to include new clustering methods and reference data sets.

## Supplementary Files

**Supplementary Information.** PDF document containing Supplementary Methods, Supplementary Results, and Supplementary Figures.

**Supplementary Table S1.** Spreadsheet file containing Supplementary Table S1.

## Acknowledgments

We thank Charlotte Soneson, Stéphane Chevrier, Vito Zanotelli, Bernd Bodenmiller, Felix Hartmann, Nikolay Samusik; and members of the Robinson, Baudis, and von Mering labs at the University of Zurich for helpful discussions and feedback on earlier drafts of the manuscript.

## References

1. Mair F, Hartmann FJ, Mrdjen D, Tosevski V, Krieg C, Becher B. The end of gating? An introduction to automated analysis of high dimensional cytometry data. European Journal of Immunology. 2016;46:34–43.

2. Saeys Y, Van Gassen S, Lambrecht BN. Computational flow cytometry: helping to make sense of high-dimensional immunology data. Nature Reviews Immunology. 2016;p. 1–14.

3. Bendall SC, Nolan GP, Roederer M, Chattopadhyay PK. A deep profiler’s guide to cytometry. Trends in Immunology. 2012;33(7):323–332.

4. Bendall SC, Simonds EF, Qiu P, Amir EaD, Krutzik PO, Finck R, et al. Single-cell mass cytometry of differential immune and drug responses across a human hematopoietic continuum. Science. 2011;332:687–696.

5. Finak G, Frelinger J, Jiang W, Newell EW, Ramey J, Davis MM, et al. OpenCyto: An Open Source Infrastructure for Scalable, Robust, Reproducible, and Automated, End-to-End Flow Cytometry Data Analysis. PLoS Computational Biology. 2014;10(8):e1003806.

6. Finak G, Langweiler M, Jaimes M, Malek M, Taghiyar J, Korin Y, et al. Standardizing Flow Cytometry Immunophenotyping Analysis from the Human ImmunoPhenotyping Consortium. Nature Scientific Reports. 2016;6:20686.

7. Aghaeepour N, Finak G, Hoos H, Mosmann TR, Brinkman R, Gottardo R, et al. Critical assessment of automated flow cytometry data analysis techniques. Nature Methods. 2013;10(3):228–238.

8. Aghaeepour N, Chattopadhyay P, Chikina M, Dhaene T, Van Gassen S, Kursa M, et al. A Benchmark for Evaluation of Algorithms for Identification of Cellular Correlates of Clinical Outcomes. Cytometry Part A. 2016;89A:16–21.

9. Hartmann FJ, Bernard-Valnet R, Quériault C, Mrdjen D, Weber LM, Galli E, et al. High-dimensional single-cell analysis reveals the immune signature of narcolepsy. Journal of Experimental Medicine. 2016;.

10. van Unen V, Li N, Molendijk I, Temurhan M, Höllt T, van der Meulen-de Jong AE, et al. Mass Cytometry of the Human Mucosal Immune System Identifies Tissue and Disease-Associated Immune Subsets. Immunity. 2016;44:1–13.

11. Pejoski D, Tchitchek N, Pozo AR, Elhmouzi-Younes J, Yousfi-Bogniaho R, Rogez Kreuz C, et al. Identification of Vaccine-Altered Circulating B Cell Phenotypes Using Mass Cytometry and a Two-Step Clustering Analysis. The Journal of Immunology. 2016;p. 4814–4831.

12. Ronan T, Qi Z, Naegle KM. Avoiding common pitfalls when clustering biological data. Science Signaling. 2016;9(432):re6.

13. Kriegel HP, Kröger P, Zimek A. Clustering High-Dimensional data: A Survey on Subspace Clustering, Pattern-Based Clustering, and Correlation Clustering. ACM Transactions on Knowledge Discovery from Data. 2009;3(1):1–58.

14. Newell EW, Cheng Y. Mass cytometry: blessed with the curse of dimensionality. Nature Immunology. 2016;17(8):890–895.

15. Chester C, Maecker HT. Algorithmic Tools for Mining High-Dimensional Cytometry Data. Journal of Immunology. 2015;195:773–779.

16. Diggins KE, Ferrell PBJ, Irish JM. Methods for discovery and characterization of cell subsets in high dimensional mass cytometry data. Methods. 2015;82:55–63.

17. Samusik N, Good Z, Spitzer MH, Davis KL, Nolan GP. Automated mapping of phenotype space with single-cell data. Nature Methods. 2016;p. 1–4.

18. Levine JH, Simonds EF, Bendall SC, Davis KL, Amir EaD, Tadmor MD, et al. Data Driven Phenotypic Dissection of AML Reveals Progenitor-like Cells that Correlate with Prognosis. Cell. 2015;162:184–197.

19. Wiwie C, Baumbach J, Ro¨ttger R. Comparing the performance of biomedical clustering methods. Nature Methods. 2015;12(11):1033–1038.

20. Hornik K. A CLUE for CLUster Ensembles. Journal of Statistical Software. 2005;14(12).

21. Spidlen J, Breuer K, Rosenberg C, Kotecha N, Brinkman RR. FlowRepository: A resource of annotated flow cytometry datasets associated with peer-reviewed publications. Cytometry Part A. 2012;81A:727–731.

22. Shekhar K, Brodin P, Davis MM, Chakraborty AK. Automatic Classification of Cellular Expression by Nonlinear Stochastic Embedding (ACCENSE). Proceedings of the National Academy of Sciences of the United States of America. 2014;111(1):202–207.

23. Chen J, Chen H. cytofkit. R package, version 1.4.8. 2016;.

24. Becher B, Schlitzer A, Chen J, Mair F, Sumatoh HR, Wei K, et al. High-dimensional analysis of the murine myeloid cell system. Nature Immunology. 2014;15(12):1181–1189.

25. Qian Y, Wei C, Lee FEH, Campbell J, Halliley J, Lee Ja, et al. Elucidation of seventeen human peripheral blood B-cell subsets and quantification of the tetanus response using a density-based method for the automated identification of cell populations in multidimensional flow cytometry data. Cytometry Part B Clinical Cytometry. 2010;78B(Suppl. 1):S69–S82.

26. Lo K, Hahne F, Brinkman RR, Gottardo R. flowClust: a Bioconductor package for automated gating of flow cytometry data. BMC Bioinformatics. 2009;10(145).

27. Aghaeepour N, Nikolic R, Hoos HH, Brinkman RR. Rapid cell population identification in flow cytometry data. Cytometry Part A. 2011;79A:6–13.

28. Finak G, Bashashati A, Brinkman R, Gottardo R. Merging mixture components for cell population identification in flow cytometry. Advances in Bioinformatics. 2009;p. 247646.

29. Ge Y, Sealfon SC. flowPeaks: a fast unsupervised clustering for flow cytometry data via K-means and density peak finding. Bioinformatics. 2012;28(15):2052–2058.

30. Van Gassen S, Callebaut B, Van Helden MJ, Lambrecht BN, Demeester P, Dhaene T, et al. FlowSOM: Using self-organizing maps for visualization and interpretation of cytometry data. Cytometry Part A. 2015;87A:636–645.

31. Sörensen T, Baumgart S, Durek P, Grützkau A, Häupl T. immunoClust - An automated analysis pipeline for the identification of immunophenotypic signatures in high-dimensional cytometric datasets. Cytometry Part A. 2015;87A:603–615.

32. Linderman M. Rclusterpp: Linkable C++ clustering. R package version 0.2.3.; 2013.

33. Zare H, Shooshtari P, Gupta A, Brinkman RR. Data reduction for spectral clustering to analyze high throughput flow cytometry data. BMC Bioinformatics. 2010;111:403.

34. Qiu P, Simonds EF, Bendall SC, Gibbs KD, Bruggner RV, Linderman MD, et al. Extracting a cellular hierarchy from high-dimensional cytometry data with SPADE. Nature Biotechnology. 2011;29(10):886–891.

35. Mosmann TR, Naim I, Rebhahn J, Datta S, Cavenaugh JS, Weaver JM, et al. SWIFT Scalable clustering for automated identification of rare cell populations in large, high-dimensional flow cytometry datasets, Part 2: Biological evaluation. Cytometry Part A. 2014;85A:422–433.

36. Nilsson AR, Bryder D, Pronk CJH. Frequency determination of rare populations by flow cytometry: A hematopoietic stem cell perspective. Cytometry Part A. 2013;83A:721–727.

37. Huber W, Carey VJ, Gentleman R, Anders S, Carlson M, Carvalho BS, et al. Orchestrating high-throughput genomic analysis with Bioconductor. Nature Methods. 2015;12(2):115–121.

